# Palaeoproteomic identification of a whale bone tool from Bronze Age Heiloo, the Netherlands

**DOI:** 10.1101/2024.04.15.589626

**Authors:** Joannes A. A. Dekker, Dorothea Mylopotamitaki, Annemieke Verbaas, Virginie Sinet-Mathiot, Samantha Presslee, Morgan L. McCarthy, Morten Tange Olsen, Jesper V. Olsen, Youri van den Hurk, Joris Brattinga, Frido Welker

## Abstract

Identification of the taxonomic origin of bone tools is an important, but often complicated, component of studying past societies. The species used for bone tool production provide insight into what species were exploited, potentially how, and for what purpose. Additionally, the choice of species may have important implications for the place of the tool within the larger toolkit. However, the taxonomic identification of bone tools is often unsuccessful based on morphology. Here we apply three palaeoproteomic techniques, ZooMS, SPIN-like data analysis and a targeted database search to narrow down the taxonomic identification of an unusually large Bronze Age bone tool from Heiloo, the Netherlands, to the North Atlantic right whale (*Eubalaena glacialis*). Additionally, the tool was investigated for use-wear, which showed that it was likely used for the processing of plant fibres. The assignment of the tool as whale bone adds support to the exploitation of whales by coastal Bronze Age populations, not just for meat, as previously suggested, but also for bone as a resource for tool production. We know of no other parallel of a bone tool such as this in terms of size, use, hafting, and taxonomic identity.

## Introduction

A common framework for studying archaeological tools and their place in past societies is to reconstruct an artefact biography, which contains the conception, manufacture, use, potential reuse and deposition of the artefact. The biography approach is commonly applied to a large variety of artefacts, but in the case of osseous artefacts the first step, the selection of raw material, remains poorly explored. The difficulty of identifying the species of a bone tool based on morphological characteristics is largely to blame for this, since morphologically diagnostic features are often removed during the shaping and use of a bone artefact.

Several biomolecular techniques have been proposed to resolve this problem, such as general applications of ancient DNA (Hofreiter and Pacher 1997; Essel et al. 2023) and shotgun proteomic analysis (Multari et al. 2023), as well as some more specialised protein-based techniques like ZooMS (Zooarchaeology by Mass Spectrometry) (Buckley et al. 2009) and SPIN (Species by Proteome INvestigation) (Rüther et al. 2022). Each of these techniques comes with their own specific advantages and disadvantages. Ancient DNA analysis is understood to be the most precise in its taxonomic identifications, but it requires a relatively high standard of biomolecular preservation.

Additionally, it often requires relatively large sample sizes (Parker et al. 2021; Tejero et al. 2024), which may not be desirable for rare archaeological objects. However, it must be noted that recent developments on minimally invasive DNA extraction may have removed this obstacle (Essel et al. 2023; Tejero et al. 2024). On the other side of the spectrum, ZooMS can often only assign taxon up to genus level, but is able to handle less well preserved samples. SPIN and shotgun proteomics seem to require a roughly similar level of preservation as ZooMS, as all are targeting ancient proteins, however, they should be able to provide more precise taxonomic identification, allowing a species level identification in cases where ZooMS can only provide a genus-level assignment. Conventional sample sizes for ZooMS and SPIN are relatively small, ranging from 5-20 mg (Rüther et al. 2022; Sinet-Mathiot et al. 2019; Welker et al. 2015), while for shotgun proteomics samples of 15-60 mg are more commonly taken (Cleland 2018; Procopio, Chamberlain, and Buckley 2017; Sawafuji et al. 2017). The development of minimally invasive sampling protocols for palaeoproteomic analysis, primarily for ZooMS, is more advanced than for aDNA analysis. A variety of different sampling protocols have been proposed that seem promising, although their ability to extract ancient protein appears to be context-specific (Evans et al. 2023; Hansen et al. 2024; McGrath et al. 2019). In this study we chose to sample destructively for three reasons. First of all, the biomolecular preservation at the site was unknown. Secondly, the size of the object allowed for a sufficient sample to be extracted without compromising other analyses of the artefact, including visual ones. Moreover, use-wear analysis was performed before destructive sampling to prevent any interference following. The combination of use-wear analysis and proteomic analysis is becoming more common and should be considered as a ‘best practice’ (Bradfield et al. 2019; Dekker et al. 2021; Orłowska et al. 2023; Hansen et al. 2024). Considering that even some minimally invasive sampling protocols for ZooMS may obscure traces of use it is vital that use-wear analysis is performed in advance to prevent the loss of valuable information regarding the use of the tool (Sinet-Mathiot et al. 2021; Hansen et al. 2024). Lastly, the aim of this research was to refine the taxonomic identification to species level for which maximising sequence coverage is essential. It has been shown that destructive methods extract a larger abundance of protein than minimally invasive methods (Hansen et al. 2024).

Palaeoproteomic techniques, both minimally invasive and destructive, have been applied in a number of previous studies to study species selection in the production of osseous tools. These studies demonstrate how palaeoproteomic techniques can be used to reveal the use of unexpected species for bone tool production (Surovell et al. 2024), such as human bone (McGrath et al. 2019; Dekker et al. 2021), as well as the intentional selection of certain species (Desmond et al. 2018; Martisius et al. 2020; Adamczak et al. 2021; Bradfield, Kitchener, and Buckley 2021).

In this study we apply multiple biomolecular techniques to a worked bone artefact (Specimen 2199) from the Bronze Age site of Heiloo-Zuiderloo, the Netherlands. Initial morphological inspection suggested that the bone artefact was of Elephantidae origin, which considering the lack of any Elephantidae species living in the Netherlands during the Bronze Age, would suggest either the use of (sub-)fossil mammoth bone or long-distance trade. To test this hypothesis we first analysed the bone artefact using ZooMS. Additionally, use-wear analysis was performed on the bone tool to study its production and use for a better understanding of the place of this artefact in the Bronze Age toolkit. However, ZooMS was not able to provide a species level identification for the bone tool, so another recently introduced palaeoproteomic technique, SPIN (Rüther et al. 2022), was performed in order to refine the taxonomic identification. The SPIN analysis revealed no collagen sequences were available for several of the species of interest. Thus, in order to resolve this blind spot, we acquired new collagen reference DNA sequences from selected cetacean species and performed a final targeted database search for a more precise taxonomic identification.

## Materials and methods

### Description of the site and context

The main subject of this study is a large plano-convex bone artefact (Specimen 2199). It was found in 2020 during archaeological excavations in Heiloo-Zuiderloo, The Netherlands (Figure 1). The site is located on the Dutch coastal area in the northwestern part of the country. The first traces of habitation in the area date from the transition period of the Neolithic to the Early Bronze Age (ca. 2000 BC). The preservation conditions of the prehistoric archaeological remains from the region are excellent due to a thick 2 metre layer of sand covering them. Additionally, both the archaeological remains as the sand layer are located below the groundwater level. As a result, many remarkable archaeological finds have been recovered in the area in recent years (Heiden 2018; de Koning and Tuinman 2019; Moesker, et al. Tol 2021; Brattinga 2023).

**Figure 1.**
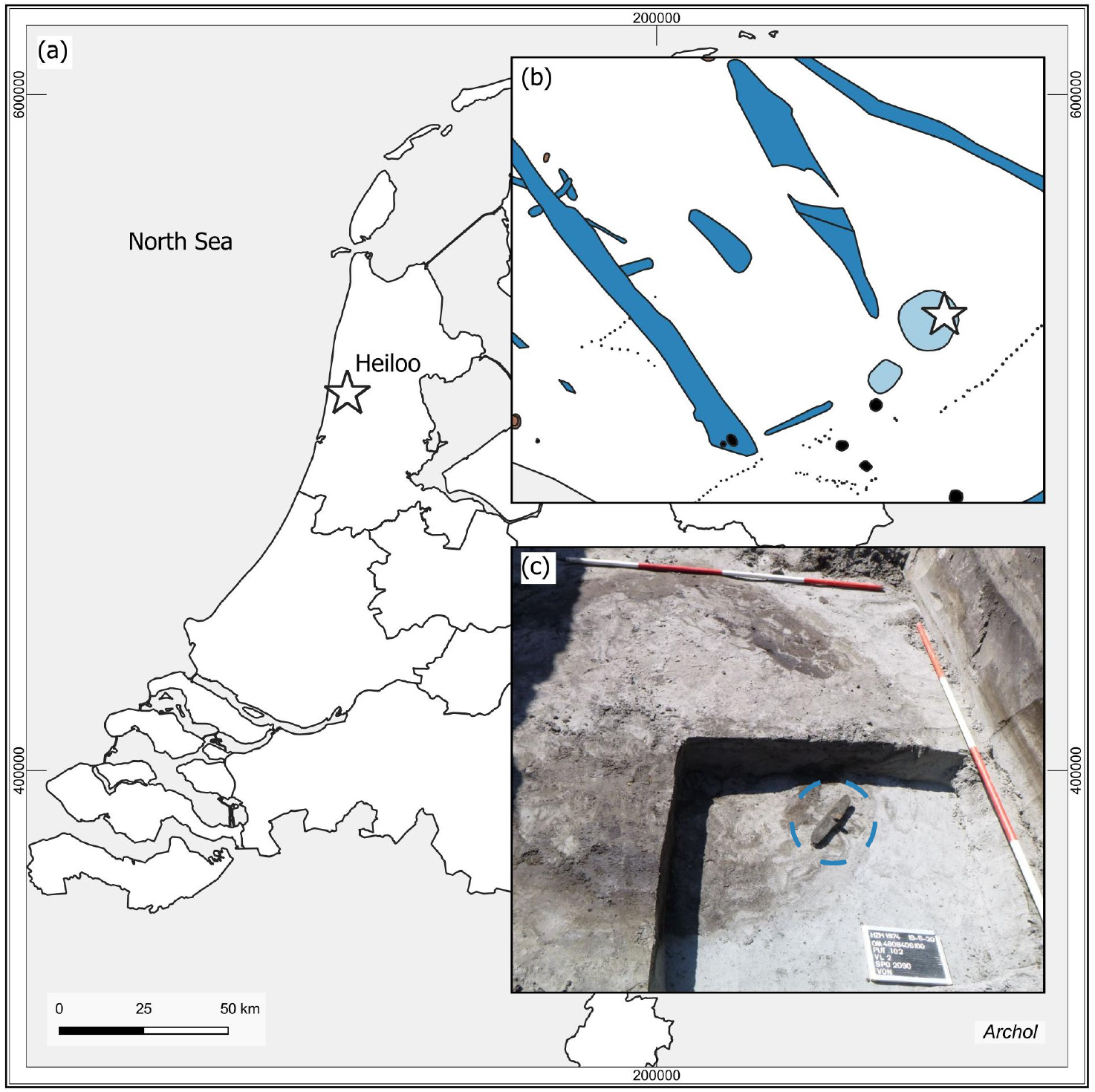
a: Map of the Netherlands with a star indicating the site location. b: Part of the excavation plan. The find location of the analysed object is marked with a star. c: Photograph showing the bone tool in situ inside the round feature of the watering hole, as indicated by the blue circle. © Archol, Leiden.

During the 2020 excavations, a Bronze Age settlement along with adjacent agricultural fields were uncovered. These findings have improved our knowledge of the Bronze Age habitation of the region, showing that the settlements were located on higher parts of the coastal landscape, specifically the dunes, while the lower parts of the landscape were utilised for various activities such as agriculture, hunting, fishing, and livestock grazing. The faunal assemblage at the site is dominated by cattle (*Bos taurus*) and sheep/goat (*Ovis aries/Capra hircus*). Other domesticates such as pig (*Sus scrofa domesticus*), dog (*Canis familiaris*) and horse (*Equus caballus*) are present in smaller numbers, as are a number of wild taxa, such as whale (Cetacea), grey seal (*Halichoerus grypus*), red deer (*Cervus elaphus*) and the common murre (*Uria aalge*). However, each of these aforementioned taxa is represented by only one or two specimens. The site also yielded a number of fish remains, the majority of which was ascribed to cod (*Gadus morhua*). The bone assemblage at the site also features a small number of worked bone artefacts. Apart from the bone tool analysed in this study, two modified cattle scapulae were found, which have been interpreted as digging implements (Nieweg 2023).

The bone object was found at the bottom of a Middle Bronze Age watering hole.

Radiocarbon dates of the wooden shaft and botanical remains from the watering hole indicate the tool was made and used around 1500 BC (Laboratory ID: Poz-125900, 3230 ± 30 BP, 1541-1425 cal. BC, laboratory ID: Poz-156692, 3290 ± 35 BP, 1631-1456 cal BC, chronological ages calculated through OxCal 4.4.4 using the INTCAL20 calibration curve (Reimer et al. 2020)).

### Description of the artefact (Specimen 2199)

The artefact has a size of 33 x 6 x 3.5 cm and has a plano-convex shape with adze-like shaped ends (Figure 2). In the middle of the object, there is a square hole measuring 2 x 2 cm where a wooden handle has been inserted at an oblique angle. The inside shape of the hole causes the shaft to be fixed with a 50 degree angle. At the thickest point, the bone is 3.5 cm thick. The upper part has a smooth and shiny surface. The other side has an irregular texture with traces of the original spongious structure of the bone. The surface of the object shows elaborate use-wear traces.

**Figure 2.**
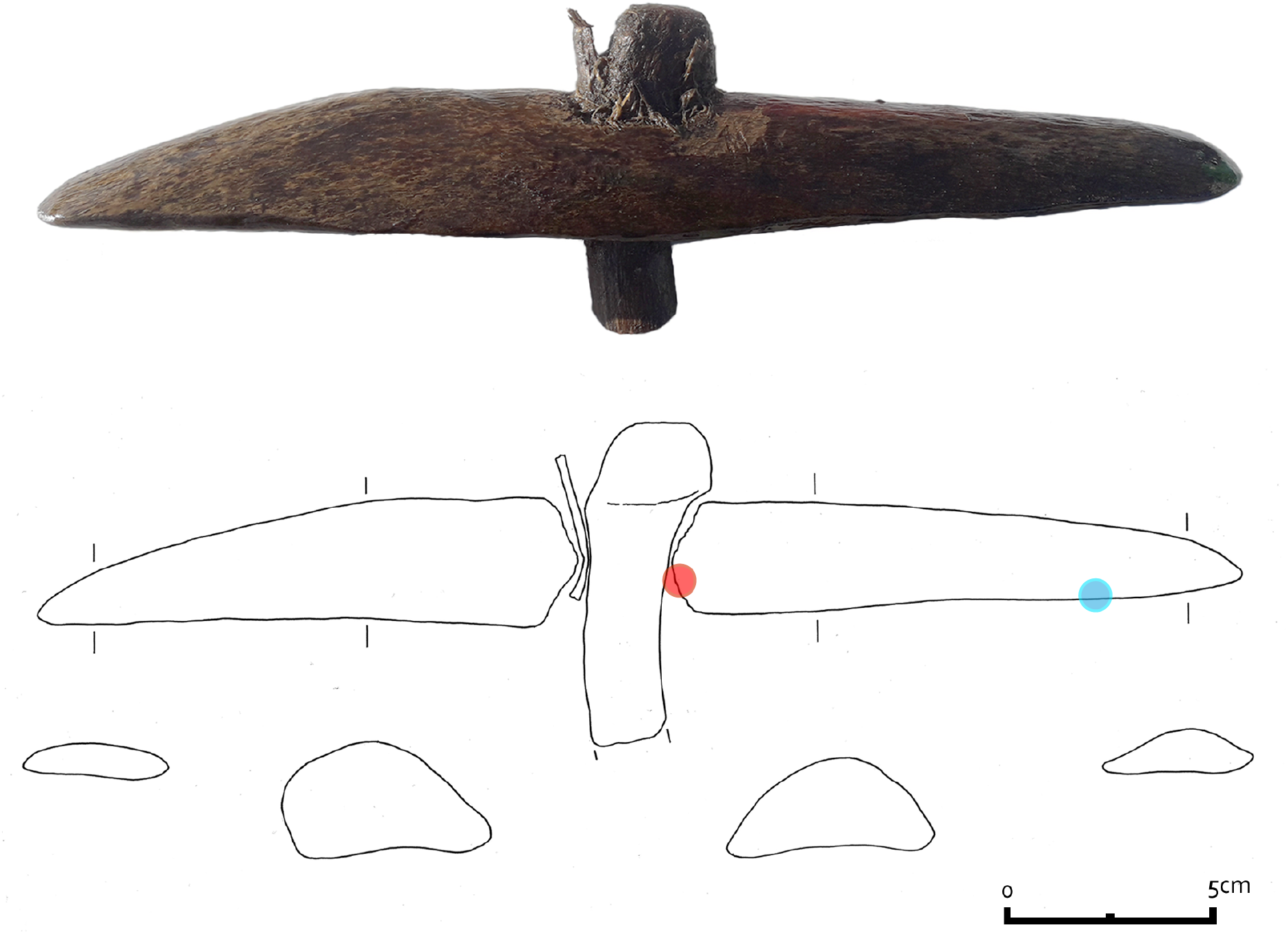
Photograph of the bone artefact, 2199, and drawing show the cross sections of the tool. The location of the first two samples is indicated by a red circle, the location of the third by a blue circle. © Archol, Leiden.

The wooden handle was damaged during excavation, resulting in the distal part of it being lost. Therefore, the original length of the handle is unknown. After the artefact was recovered, it became clear that the handle was secured within the shaft of the tool by a wooden wedge. The wedge is inserted between the bone and the handle from the front and is made from the outer part of a willow branch. Based on the anatomical features of the cross, radial and tangential section it was determined that the handle itself is made from a modified oak branch (van Hees and van Amerongen 2023). The branch is thickened 2 cm from the top, where a knob has been shaped using a sharp metal object. The finishing marks on the knob are relatively coarse and functional.

Three samples consisting of small bone flakes (28.6 mg, 21.0 mg and 10.7 mg) were taken from the artefact by scraping the bone with a scalpel. Two were taken from the inside of the shaft hole, the third from the downside of the artefact. The first sample was sent to BioArCh, University of York, the second to the Department for Evolutionary Anthropology of the Max Planck Institute in Leipzig, and the third sample was sent to the Globe Institute, University of Copenhagen. The sample to York was designated for ZooMS analysis, whereas the sample for Leipzig was analysed with both ZooMS and LC-MS/MS. Lastly the sample sent to Copenhagen would be analysed using LC-MS/MS. ZooMS analysis was performed at York and Leipzig. Although unintended, an independent validation of the obtained MALDI-ToF MS spectra provides further validation of the resulting taxonomic identification of this unique bone artefact. From our knowledge, inter-laboratory comparisons are either rare in ZooMS, or have so far remained unreported. Although we do not conduct a formal inter-laboratory comparison either, we do argue that periodic inter-laboratory tests, even on small scales, may be beneficial. In particular, with a larger variety of MALDI-ToF MS instrumentation starting to be used by the ZooMS community, such comparisons might shed light on the benefits of particular mass spectrometry application for particular ZooMS applications.

### Use-wear methods

Apart from the biomolecular analysis, use-wear analysis was also undertaken in order to investigate the function of the bone tool. Use-wear analysis is based on a visual comparison of the traces on archaeological tools and experimental replicas. The method has been commonly used in archaeology during the last decade (e.g. le Moine 1994, Van Gijn 2007 and Evora 2015). For the analysis of the object described here a Leica M80 stereomicroscope (magnifications 7.5 - 60 x) and a Leica DM2700 metallographic microscope (magnifications 100 and 200 x) and the reference collection of the Leiden University laboratory for Material Culture studies were used. Photos were taken with a Leica MC120HD camera.

### ZooMS

Peptides were extracted for ZooMS analysis using both the cold acid and the ammonium bicarbonate (AmBic) extraction protocols (van Doorn, Hollund, and Collins 2011). The samples at both York and Leipzig were treated according to similar protocols. In short, the cold acid protocol consists of demineralising the bone fragment in 0.6 M hydrochloric acid (HCl) at 4°C for 18 hours, followed by the removal of the acid supernatant and neutralisation using 50 mM AmBic (NH_4_HCO_3_, pH 8.0). The proteins in the samples are then gelatinised via an incubation in 50 mM AmBic at 65°C for one hour. Subsequently, 50 µL of the resulting supernatant was digested using trypsin (Promega) at 37°C, acidified using trifluoroacetic acid (10% TFA), and then cleaned and desalted on C18 ZipTips (Thermo Scientific). Lastly, the filtered peptides eluted in 0.1% TFA in 50:50 acetonitrile and UHQ water are spotted in triplicate on a MALDI plate along with α-cyano-4-hydroxycinnamic acid (CHCA, Sigma) as a matrix solution. The plate was then analysed using an autoflex LRF MALDI-TOF (Bruker) in Leipzig in reflector mode, positive polarity, and matrix suppression up to 590 Da and collected in the mass-to-charge range 1000–3500 m/z; and a Ultraflex III MALDI-TOF (Bruker) in York in reflector mode, positive polarity, peptide masses below 650 were suppressed and the mass range was set between 800-4000 m/z . Before sample acquisition, external mass calibration was achieved on adjacent MS standard spots with a peptide calibration standard (In Leipzig: #8206195, Bruker Daltonics, Germany) containing a mixture of seven peptides (Angiotensin II m/z = 1046.541, Angiotensin I m/z = 1296.685, Substance_P m/z = 1347.735, Bombesin m/z =1619.822, ACTH (1–17 clip) m/z = 2093.086, ACTH (18–39 clip) m/z = 2465.198 and Somatostatin m/z = 3147.471. In York: Des-Arg1 Bradykinn m/z = 904.681, Angiotensin I m/z = 1295.685, Glu1-Fibrino- peptide B m/z = 1750.677, ACTH (1–17 clip) m/z = 2093.086, ACTH (18–39 clip) m/z = 2465.198 and ACTH (7–38 clip) m/z = 3657.929).

The AmBic protocol differs from the cold acid protocol by skipping the acid demineralisation step. Instead the bone is immediately incubated in 50 mM AmBic at 65°C for one hour. Afterwards the AmBic protocol is the same as the cold acid protocol in terms of digestion, peptide filtering and mass spectrometry. Consequently, the main difference between the two methods is that the cold acid protocol targets the acid-insoluble collagen fraction of the bone, while the AmBic protocol focuses on the AmBic-soluble component of the collagen.

The obtained mass spectra were matched against the curated database of ZooMS biomarkers from the University of York (Presslee 2020), as well as specific publications on cetacean biomarkers (Buckley et al. 2009, 2014; Kirby et al. 2013; Speller et al. 2016; Welker et al. 2016). The obtained raw spectra can be found at Zenodo under the identifier 10970629. The observed biomarkers allowed us to assign a family level identification to specimen 2199. In order to further specify the taxonomic identification, it was decided to continue with shotgun-proteomic analysis for this specimen.

### LC-MS/MS data acquisition for SPIN data analysis

The “Species by Proteome INvestigation” (SPIN) approach is a recently proposed proteomics workflow leveraging automatic approaches to LC-MS/MS data analysis in association with shorter liquid chromatography separation and DIA or DDA spectral acquisition (Rüther et al. 2022). The workflow employed in this study uses the bioinformatic tools published in SPIN, but follows a different protein extraction protocol. Samples were first processed using the AmBic protocol as outlined above, then demineralized and further processed as the cold acid protocol outlined above, save for a few minor modifications after trypsin digestion. This follows previous strategies employed in ZooMS studies (Welker et al. 2016) and recommendations made for DDA-based SPIN analysis (Mylopotamitaki et al. 2023). After digestion, peptides for each extraction protocol were combined by sample location and loaded onto a single StageTip (Rappsilber et al. 2007) together, prior to mass spectrometry analysis. Peptides were eluted from the StageTips with 30 µL of 40% acetonitrile (ACN) and 0.1% formic acid (FA) and vacuum centrifuged until less than 3 µL remained. The samples were then resuspended in 8 μL of 0.1% trifluoroacetic acid (TFA), 5% ACN. Of each sample 0.5 µL was loaded on an EASY-nLC 1200 (Thermo Fisher Scientific) connected to an Orbitrap Exploris 480 (Thermo Fisher Scientific). Mass spectrometric data was acquired using previously published parameters for archaeological samples (Brandt et al. 2023). A separate blank extraction was performed alongside the adze samples, resulting in a total of three LC-MS/MS injections. The resulting proteomics data was deposited to the ProteomeXchange Consortium via PRIDE (Perez-Riverol et al. 2022) with the dataset identifier PXD051408.

### MaxQuant data analysis

Two MaxQuant searches were performed. For the first, the three .raw files were analysed in MaxQuant version 1.6.7.0 against a custom database containing concatenations of the mature COL1A1 and COL1A2 sequences of 54 mammalian species, i.e. the triple helical region and the telopeptides. The database includes 13 cetaceans, of which 8 are protein sequences derived from the genomic resources provided by (Árnason et al. 2018). Non-cetacean sequences were downloaded from NCBI. Variable modifications that were included in the search, were oxidation (M), deamidation (NQ), Gln -> pyro-Glu, Glu -> pyro-Glu, and proline hydroxylation, with protease specificity set to Trypsin/P in specific cleavage mode.

The second search was performed with MaxQuant version 2.4.10.0 with the same three .raw files, but against a more constrained database (SI 1), consisting of the COL1A1 and COL1A2 sequences of 14 and 12 baleen whale species, respectively. These collagen sequences were acquired from publicly available resources and additional sequences obtained by translating protein sequences for baleen whale genomic resources available online (see Bioinformatics data analysis section below). For this second search oxidation (M), acetyl (protein N-term), deamidation (NQ) and proline hydroxylation were selected as variable modifications. A maximum number of six modifications per peptide was allowed and the protease specificity was set to trypsin/P. A maximum of two missed cleavages was allowed for each peptide. Allowing semi-specific trypsin cleavage can increase the number of detected peptides, especially in more degraded samples where protein fragments are relatively short already before digestion. The enzymatic digestion of these shorter protein fragments creates a relatively large amount of peptides with a semi-specific cleavage pattern, because the other end of the peptide had been broken due to diagenesis rather than enzymatic digestion. Allowing semi-tryptic cleavage in the protein analysis will facilitate the identification of such peptides, but this is not always the case. The increase in search space caused by the inclusion of semi-tryptic peptides may inflate the number of decoy matches, which in turn can result in a reduced number of target peptide matches as a higher score threshold must be used to maintain the same false discovery rate (FDR) (Fahrner et al. 2021; Palomo et al. 2023). We chose to only search for fully tryptic peptides as the initial results indicated relatively good preservation.

### Bioinformatics data analysis

#### Genomic protein translation

Raw reads from 23 baleen whales, representing 15 different species, were accessed from the European Nucleotide Archive or NCBI Sequence Read Archive (SI 2). As genetic data was missing for the Bryde’s whale (*Balaenoptera brydei*), we generated a 30x genome from a juvenile male (specimen MCE1246) stranded on September 1st 2000 in Isefjorden, Denmark. DNA was extracted using the Thermo Scientific KingFisher Cell and Tissue DNA Kit and sequenced on a DNBSEQ at BGI China. Accessed reads were trimmed using the bbduk.sh script from bbmap (Bushnell 2014), with the settings ktrim=r, k=23, mink=8, hdist=1, tbo, qtrim=rl, trimq=15, maq=20, and minlen=40. The blue whale (*Balaenoptera musculus*) reference genome (Bukhman et al. 2022) was selected for read mapping given its status as a platinum-standard reference genome (Morin et al. 2020). Trimmed reads were mapped using the Burrows-Wheeler Aligner Maximal Exact Matches (BWA-MEM) algorithm v.0.7.17 and default settings (Li and Durbin 2009). PCR duplicates were removed with SAMtools v.1.17 (Li et al. 2009). The target protein transcript sequences were extracted from the mapped (bam) files with the SAMtools view command and genomic coordinates from the reference genome annotations (COL1A1 - CM020960.2: 24504018 - 24520329; COL1A2 - CM020949.2: 58946669 - 58982479). Consensus sequences were generated from the bam files using ANGSD v.0.940-2 (Korneliussen et al. 2014) with the parameters doFasta 2, doCounts 1, minQ 30, and minMapQ 30. To ensure high confidence in the final translated protein sequences, the following steps were manually applied to each consensus sequence individually in the Geneious Prime v.2020.2.5 (Kearse et al. 2012) graphical user interface. The annotated coding sequences for the blue whale COL1A1 (NCBI Accession ID: XM_036835733.1) and COL1A2 (NCBI Accession ID: XM_036862852.1) genes were each aligned to the corresponding consensus transcript sequences generated for each species with MAFFT v.7.490 (Katoh and Standley 2013; Katoh et al. 2002). Gaps in the alignment representing untranslated regions (UTRs) were trimmed and the resulting coding sequence from the target species was extracted and translated to the protein sequence in Geneious Prime.

To validate this approach, we compared our resulting protein sequences for those previously annotated for the blue whale reference genome COL1A1 (NCBI Accession ID: XP_036691628.1) and COL1A2 (NCBI Accession ID: XP_036718747.1). Validation also included comparing our protein sequences with the annotated protein sequences from the fin whale (*Balaenoptera physalus*) and North Atlantic right whale (*Eubalaena glacialis*) assembly websites (dnazoo.org/assemblies) generated by the DNAzoo consortium (Dudchenko et al. 2017). Through alignment with existing annotated protein sequences for the fin whale and North Atlantic right whale, we observed that the blue whale COL1A1 transcript variant X1 (XP_036691628.1) had an additional 43 amino acids at amino acid positions 433 to 475. This protein sequence region was trimmed from each of our translated protein sequences for consistency with publicly available annotations. Two accessions (SRR13167951; *Eubalaena australis* (southern right whale) and SRR935201; fin whale (*Balaenoptera physalus*), yielded consensus sequences with excessive missing bases and were excluded from the analysis. In addition to the curated sequences, we concatenated 12 publicly available sequences (SI 1) to comprise a final COL1 dataset of 33 baleen whales, representing 14 species.

#### Mass spectrometric data

Subsequent to spectral identification, data analysis was conducted largely through R (version 4.1.2) in RStudio (version 2022.02.0.0) using the packages tidyverse (version 1.3.1) (Wickham et al. 2019) seqinr (version 4.2-8), ggpubr (version 0.4.0), data.table (version 1.14.2) (Barett et al. 2024), bit64 (version 4.0.5) (Karunarathne et al. 2022), ggsci (version 2.9), progressr (version 0.10.0) (Bengtsson 2023), and stringi (version 1.7.6), MALDIquant (version 1.22.2) (Gibb and Strimmer 2012), MALDIquantForeign (version 0.14.1) (Gibb 2024), gridExrta (version 2.3) (Auguie 2017). For SPIN data analysis, the peptides detected in the first MaxQuant search (see above) were analysed using the DDA script with default settings and protein sequence database provided by Rüther et al. 2022. This protein database of 20 bone proteins across 156 mammalian species contains 15 cetacean species of which 4 are baleen whale entries. After ZooMS and SPIN, and having determined that the bone artefact probably derives from a baleen whale, we conducted subsequent MaxQuant and amino acid sequence analysis against a sequence alignment of 29 baleen whale sequences, representing 14 species for COL1A1 and 25 sequences, representing 12 species for COL1A2. An overview of which whale genomes provided reference for COL1A1 and/or COL1A2 can be found in SI 2. We determined all relevant single amino acid polymorphisms (SAPs) at which species of the baleen whale genera *Balaena* and *Eubalaena* differ, and compared these SAPs with the coverage obtained in the second MaxQuant database search.

Finally, the deamidation rate of the proteins observed during the targeted MaxQuant search, excluding any contaminants, was calculated using the following Mackie et al. 2018.

## Results

### Use-wear results

Inspection of the bone artefact shows that the tool was formed from a longitudinal section of the bone, as the trabecular bone is visible. According to the observed use-wear traces, the first shaping was done by cutting, chopping and grinding (Figure 3A). Exactly how the rounded ends were shaped is unclear, as any traces of this were removed by subsequent grinding to give them their final shape. The hole is square in shape and was made using a chisel (Figure 3B), working from both sides of the tool. On the inside of the hole, clear wear of a shaft is visible, indicating prolonged use. The tool was bound to the haft using a rope that extended to about midway between the hole and the edges of the tool. The traces left by the binding materials show both characteristics of plant and animal materials (Figure 3C). This can either indicate the use of bast as a binding material or the use of different binding materials in different stages of the tools use-life. This could not be interpreted with certainty. The traces on the edges of the tool are unfortunately less clear and one of the edges is damaged, which removed most of the use-wear traces. The traces that are visible are fatty, and in some spots smooth in character. The edge is rounded and there are clear striations visible in the polish (Figure 3D). This leads to the interpretation that the tool was used on a plant material. Based on the rounding of the edge it was not used to chop, but more probably the pounding or the processing of fibres in another manner.

**Figure 3.**
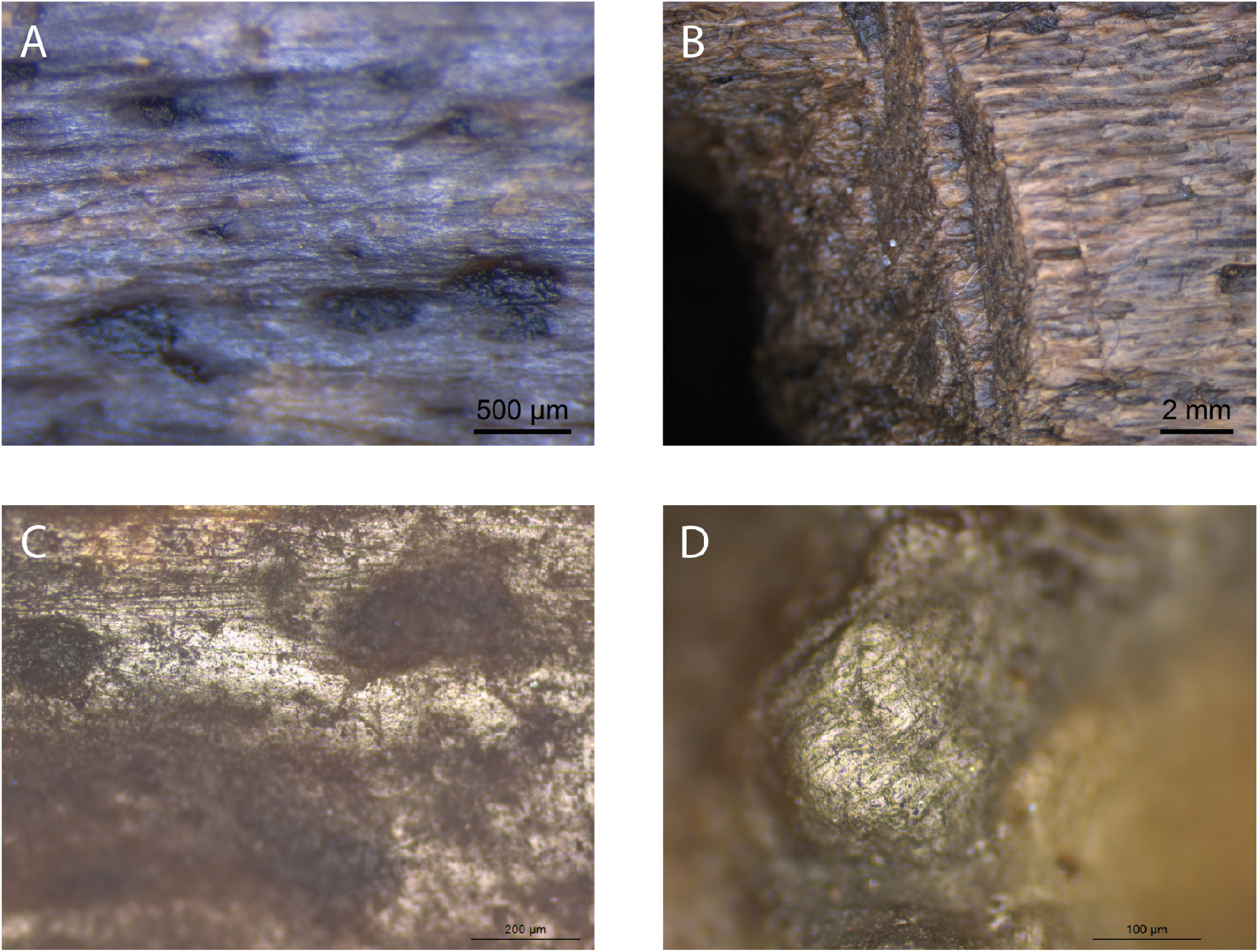
Photos of the use-wear on the bone object. A, traces of grinding on the lower part of the tool (original magnification 40x). B, traces of chiselling in the hole (original magnification 7,5x). C, traces of the binding material used to bind the tool to the haft (original magnification 100x). D, traces on the edge of the tool (original magnification 200x).

### ZooMS

ZooMS extraction was performed independently in two different locations and analysed on separate MALDI TOF-MS instruments. Three out of four extractions of the bone artefact provided consistent results attributing the sample to *Eubalaena glacialis* (North Atlantic right whale) or *Balaena mysticetus* (bowhead whale) (Table 1). The fourth sample provided inconsistent results with marker masses observed that suggest a mixture of taxonomic sources, preventing an adequate taxonomic identification through ZooMS for this extraction. Nevertheless, the consistency of the marker masses observed among the other three extracts (Figure 4), generated in two laboratories and analysed on different MALDI TOF-MS instruments, gives confidence that according to ZooMS the bone artefact likely belongs to the Balaenidae family.

**Figure 4.**
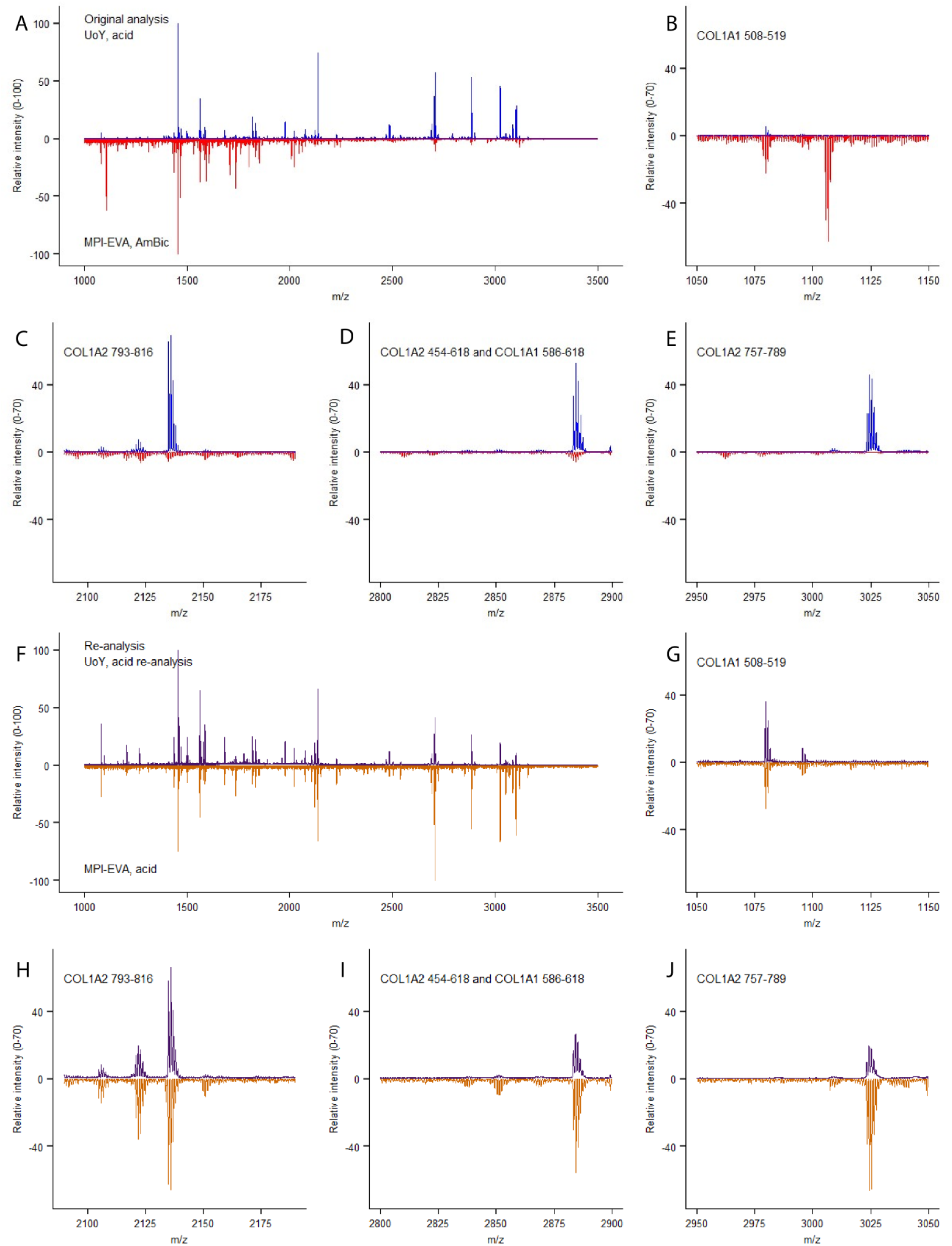
ZooMS spectra. A, total spectra of the original UoY (blue) and MPI-EVA (red) analyses. B-E, close-ups of the mass spectra for biomarkers COL1A1 508-519, COL1A2 793-816, COL1A2 454-618 AND COL1A1 586-618, COL1A2 757-789 respectively. F, total spectra of the re-analysed samples from UoY (purple) and MPI-EVA (orange). G-J, zoomed in spectra of the biomarkers COL1A1 508-519, COL1A2 793-816, COL1A2 454-618 AND COL1A1 586-618, COL1A2 757-789 respectively.

**Table 1.** Observed biomarkers and matching taxa, oxidised variants of biomarkers are notated in parentheses, *indicates that a possible peak at 1161.6 m/z was also observed, although at a lower intensity than 1189.6 m/z. UoY denotes a protein extraction performed at the University of York, whereas MPI-EVA indicates samples extracted at the Max Planck Institute, Department for Evolutionary Anthropology, Leipzig. “Acid” signifies that the sample was extracted using the cold acid extraction (van Doorn, Hollund, and Collins 2011), “AmBic” indicates an AmBic buffer extraction protocol (van Doorn, Hollund, and Collins 2011). The NA right whale refers to the North Atlantic right whale (*Eubalaena glacialis*).

| Sample | COL1a1<br>508 - 519 | COL1a2<br>978 - 990<br>/ (+16) | COL1a2<br>484 - 498 | COL1a2<br>502 - 519 | COL1a2<br>292 - 309 | COL1a2<br>793 - 816 | COL1a2<br>454 - 483 | COL1a1<br>586 - 618<br>/ (+16) | COL1a2<br>757 - 789<br>/ (+16) | Latin name | Common<br>name |
| --- | --- | --- | --- | --- | --- | --- | --- | --- | --- | --- | --- |
| UoY, acid,<br>2199 | 1079.6 | 1205.6 | 1453.7 | 1566.8 | 1682.8 | 2135.1 |  | 2883.4 /<br>2899.4 | 3007.4 /<br>3023.4 | *Eubalaena glacialis*<br>/ *Balaena*<br>*mysticetus* | NA right whale<br>/ Bowhead<br>whale |
| UoY, acid,<br>reanalysis,<br>2199 | 1079.6 | 1189.6 /<br>1205.6 | 1453.7 | 1566.8 | 1682.8 | 2135.1 | 2849.4 | 2883.4 /<br>2899.4 | 3007.4 /<br>3023.4 | *Eubalaena glacialis*<br>/ *Balaena*<br>*mysticetus* | NA right whale<br>/ Bowhead<br>whale |
| MPI-EVA,<br>AmBic,<br>2199 | 1079.6/<br>1105.6 | 1161.6 | 1453.7 | 1550.7 /<br>1566.7 | 1648.8 /<br>1682.8 | 2115.8 | 2808.3 | 2883.4 /<br>2899.4 | 2999.2 | Contaminated | - |
| MPI-EVA,<br>acid, 2199 | 1079.6 | 1189.6* /<br>1205.6 | 1453.7 | 1566.8 | 1682.8 | 2135.1 | 2849.4 | 2883.4 /<br>2899.4 | 3007.4 /<br>3023.4 | *Eubalaena glacialis*<br>/ *Balaena*<br>*mysticetus* | NA right whale<br>/ Bowhead<br>whale |

### SPIN data analysis

Since the ZooMS taxonomic assignment of the artefact specimen 2199 leaves several baleen whale species as the possible taxonomic origin, we proceeded with separate proteomic extractions for shotgun proteomic analysis. One extraction included material sampled from the inside of the hafting hole of the artefact, while the other extraction included material sampled from the outside of the artefact. The generated data was first analysed using the SPIN analysis workflow and the SPIN mammalian bone protein database, which managed to identify the amino acid sequences for 419 peptides from the outside of the bone tool and 469 peptides from the inside of the artefact. These peptides were matched to a variety of different proteins with varying degrees of taxonomic specificity. Taxonomic identifications (Table 2) were obtained by evaluating the frequencies of proteins matching a particular taxon, resulting in a SPIN assignment to *Balaeonoptera acutorostrata*. The extraction blank was matched to *Bos* sp. by the SPIN script, but it has been excluded from Table 2 as 84% of the peptide spectral matches (PSMs) in the blank were derived from the trypsin used for digestion. This also means that the number of PSMs used by the SPIN script to assign a taxonomic identification to the blank sample is very low and most likely derived from background contamination. The blank intensity signal can therefore be regarded as background noise.

**Table 2.** SPIN taxonomic identifications of the bone tool (2199). The Site count refers to the number of amino acid positions called with high confidence and present in the protein sequences of the highest-ranking species entry (here, *Balaenoptera acutorostrata*). The Non-matching site count indicates the remaining high-confidence amino acid positions that do not match the highest-ranking species entry.

| Sample | Latin name | Common name | Site count | Non-matching site count | Relative protease intensity (%) |
| --- | --- | --- | --- | --- | --- |
| 2199, inside hafting hole | *Balaenoptera acutorostrata* | Common minke whale | 2310 | 159 | 1.1 |
| 2199, lower side artefact | *Balaenoptera acutorostrata* | Common minke whale | 2392 | 169 | 1.0 |

At first glance it seems that the SPIN results, an assignment to *Balaenoptera acutorostrata*, contradict the ZooMS results, with an assignment to either *Eubalaena glacialis* or *Balaena mysticetus*. However, the SPIN database only contains entries for four closely-related baleen whale species within the genus *Balaenoptera*, whereas the ZooMS peptide marker database includes entries for eight baleen whale species, including *Eubalaena glacialis*, *Balaena mysticetus*, and *Balaenoptera acutorostrata*. This suggests the SPIN taxonomic identification might be driven by a lack of representative sequence entries in case the "true" taxonomic identity lies outside the four species included within its database.

### Selected baleen whale database search

To resolve this potential conflicting taxonomic identification, we translated COL1A1 and COL1A2 sequences from genomic resources available for 14 baleen whale species. Next we performed a second MaxQuant search against the resulting database of all available eight COL1A1 and five COL1A2 baleen whale sequences plus the additional sequences translated from the acquired genomic data for a total of 14 and 12 species for COL1A1 and COL1A2, respectively. Our COL1A1 and COL1A2 sequence database therefore contained a sequence entry for each known baleen whale genus and species, except *Balaenoptera omurai* (Omura’s whale) and *Balaenoptera edeni* (Eden’s whale, a small form of Bryde’s whale).

We reanalysed our shotgun proteomic data against this new baleen whale-specific sequence database containing COL1 entries only. The protein groups identified in this more targeted search were filtered by removing any matches to a decoy sequence, as well as those protein groups that only had two or fewer unique peptides. Additionally, we monitored the prevalence of deamidation using a publicly available Python script (Mackie et al. 2018) as a rough indicator of modern contamination. The results indicate that on average 52.7% of the asparagine residues in the bone object were deamidated, as well as 28.4% of the glutamine residues. These values fall in the overlapping range of previously published deamidation values for modern and archaeological samples (Ramsøe et al. 2020; Pal Chowdhury and Buckley 2022). However, the standard deviation of deamidation in modern bones is so large that even a sample from a 46-107 ka cave site fell within their range (Brown et al. 2021). The deamidation values therefore raise no doubts regarding the age of the extracted collagen.

These criteria left one protein group for each of the two proteins included in the database (Table 3). Both the protein groups contain collagen variants of three different species. The COL1A1 protein group contains peptides matching to the three species of the genus *Eubalaena*, while the COL1A2 protein group also includes the species *Balaena mysticetus*. Aligning the COL1A1 and COL1A2 sequences of *Eubalaena* sp. and *Balaena mysticetus* reveals five SAPs that would allow distinguishing between the two taxonomic groups. For each of the five SAPs the number of PSMs matching the *Eubalaena* sp. and the *Balaena mysticetus* reference sequence were counted to provide an overview of the support for both taxa (Table 4). The taxonomic specificity of the five SAPs was checked by searching the longest observed peptide covering each SAP against the NCBI nr database using BLAST (Altschul et al. 1990). Though it must be noted that these five peptides were not selected for being unique to either *Eubalaena glacialis* or *Balaena mysticetus*, but for differing between the two species. The ZooMS and SPIN-analysis results enable us to refine the taxonomic identification starting from the level of baleen whales. Nevertheless, it is important to check the specificity of the target peptides to see if they are shared with common contaminant taxa.

**Table 3.** Number of peptides per species for the most abundant protein group for COL1A1 and COL1A2.

| Protein | Species | Peptide count (razor + unique) |
| --- | --- | --- |
| COL1A1 | *Eubalaena australis* | 135 |
| COL1A1 | *Eubalaena japonica* | 135 |
| COL1A1 | *Eubalaena glacialis* | 135 |
| COL1A2 | *Eubalaena glacialis* | 109 |
| COL1A2 | *Eubalaena japonica* | 109 |
| COL1A2 | *Balaena mysticetus* | 109 |

**Table 4.**
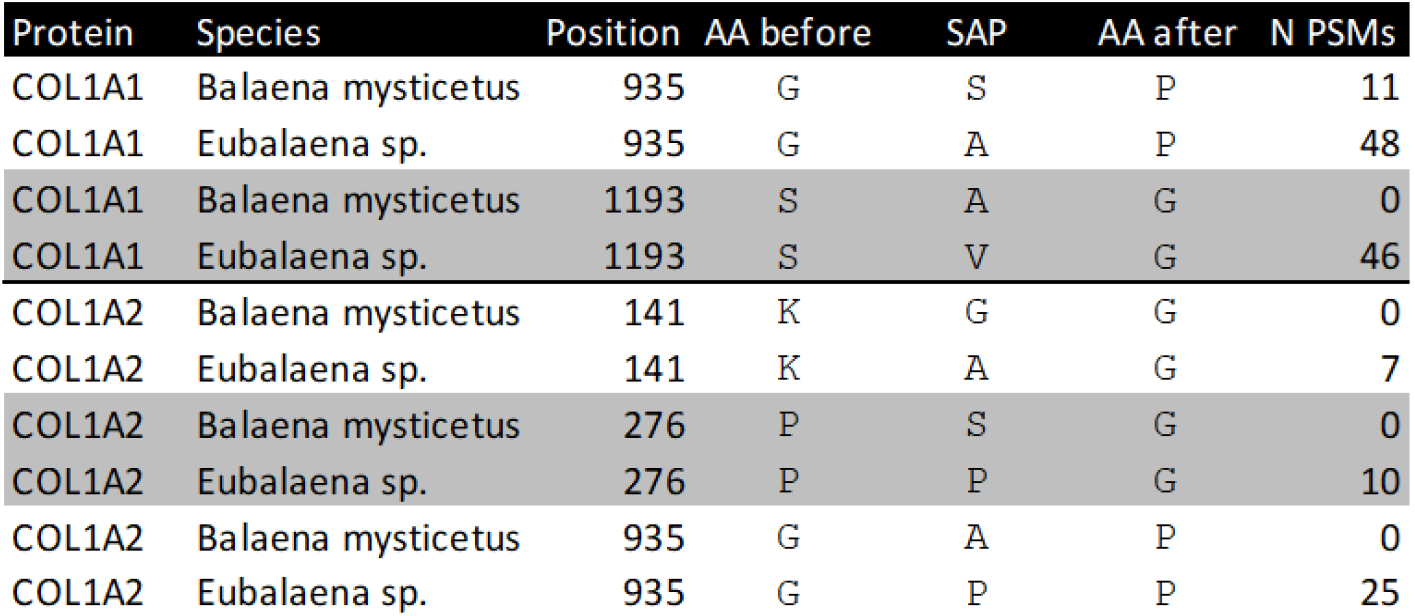
SAPs differing between *Eubalaena* sp. and *Balaena mysticetus*, and the number of matching observed PSMs.

| Protein | Species | Position | AA before | SAP | AA after | N PSMs |
| --- | --- | --- | --- | --- | --- | --- |
| COL1A1 | Balaena mysticetus | 935 | G | S | P | 11 |
| COL1A1 | Eubalaena sp. | 935 | G | A | P | 48 |
| COL1A1 | Balaena mysticetus | 1193 | S | A | G | 0 |
| COL1A1 | Eubalaena sp. | 1193 | S | V | G | 46 |
| COL1A2 | Balaena mysticetus | 141 | K | G | G | 0 |
| COL1A2 | Eubalaena sp. | 141 | K | A | G | 7 |
| COL1A2 | Balaena mysticetus | 276 | P | S | G | 0 |
| COL1A2 | Eubalaena sp. | 276 | P | P | G | 10 |
| COL1A2 | Balaena mysticetus | 935 | G | A | P | 0 |
| COL1A2 | Eubalaena sp. | 935 | G | P | P | 25 |

Of the five, the COL1A1 peptides covering the SAP at position 935 and 1193 did not match to any other species. Unfortunately, BLAST obtained no significant match for the COL1A2 peptide covering the SAP at position 141 and the peptide covering the SAP at position 276 was shared with *Balaenoptera acutorostrata, Balaenoptera ricei* and *Balaenoptera musculus*. Lastly, the COL1A2 peptide covering the SAP at position 935 was found to be shared among several other taxonomic groups. These were several species of bats (Chiroptera), three Metatherian species and 13 toothed whale species (Odontoceti). However, the only baleen whale species it matched was *Eubalaena glacialis*. It should be emphasised that although BLAST is often considered the golden standard for taxonomic identification, the NCBI nr database it relies on also suffers from incomplete references.

Consequently, a custom database, as used here, may be able to achieve more precise taxonomic identifications.

For four of the five SAPs the observed peptides only matched with the *Eubalaena* sp. sequences, but for the COL1A1 SAP at 935 there were also a number of PSMs matching the *Balaena mysticetus* sequence. To further validate the peptide identifications, the MS2 spectra and alignment of the respective peptides of the SAPs at position 935, 1193 for COL1A1 and at position 276 for COL1A2 were visualised (Figures 5-7). These show that the fragment ion coverage of the SAPs is quite extensive for all the relevant peptides and argues in favour of the validity of these peptide sequence identifications, including the contrasting sequences for the SAP at COL1A1 935. The majority of the protein evidence suggests *Eubalaena* sp. as the source of the bone artefact (Specimen 2199), as for all SAPs several peptides matching to the *Eubalaena* sp. reference were found.

**Figure 5.**
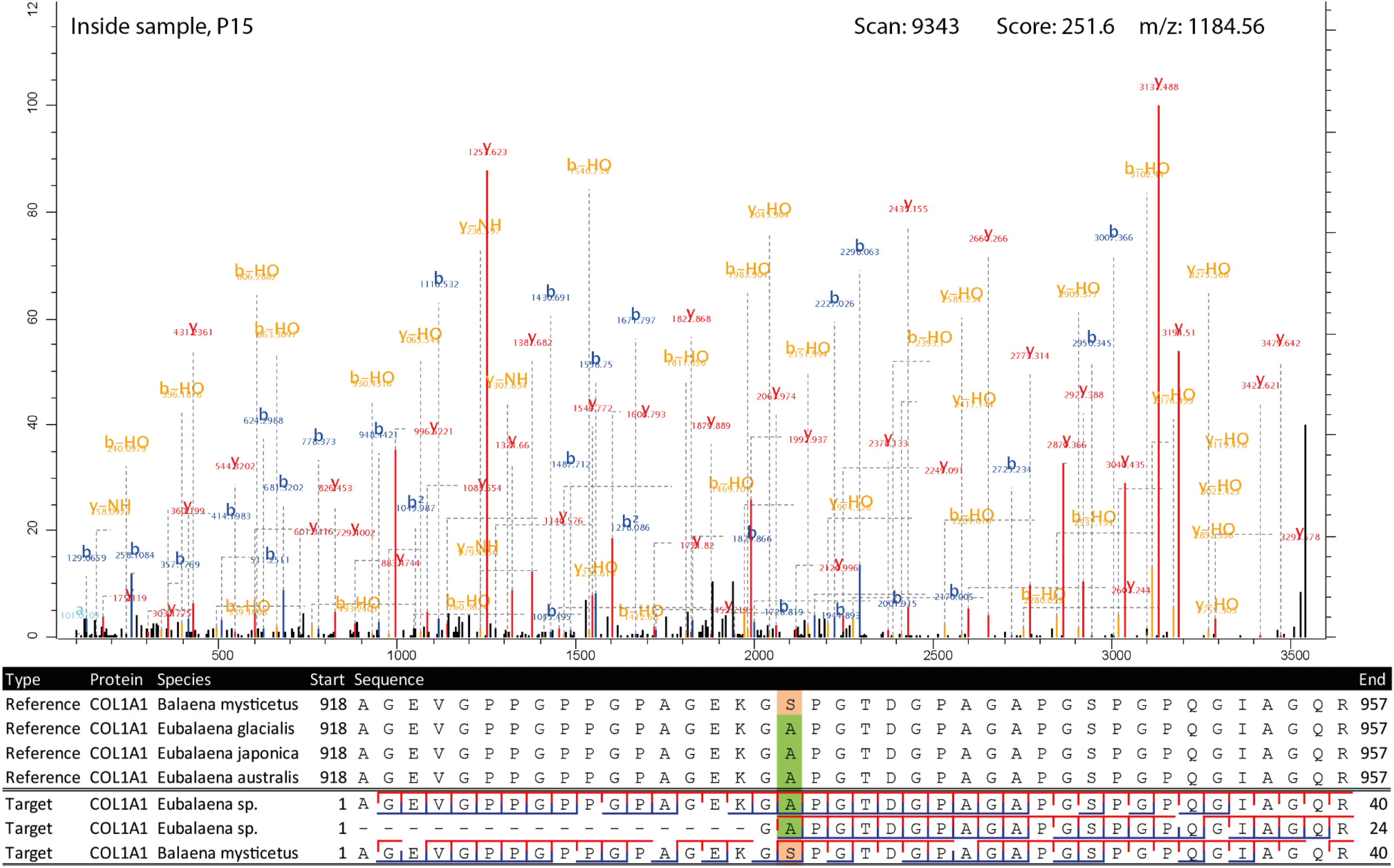
Highest scoring MS2 spectrum of the COL1A1 918-957 *Eubalaena* sp. peptide covering the 935 SAP. The spectra matching the *Eubalaena* sp. sequences are highlighted in green at the SAP, whereas the sequences matching the *Balaena mysticetus* reference are highlighted in light orange. The b-ion coverage is annotated in blue, and the y-ion coverage in red.

**Figure 6.**
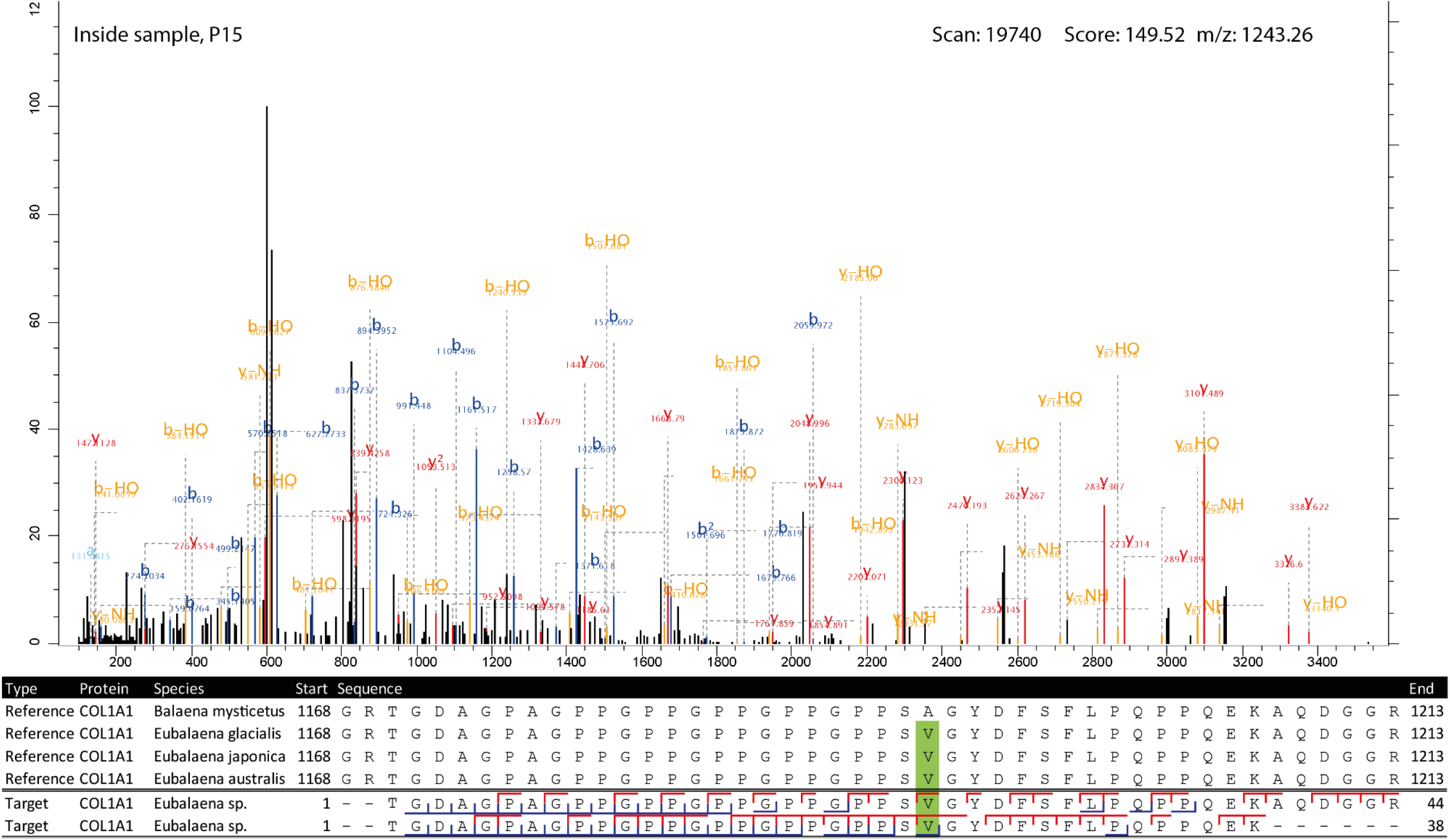
Highest scoring MS2 spectrum of the COL1A1 1170-1213 peptide covering the 1193 SAP. The reference spectra matching the observed peptides are highlighted in green at the SAP. The b-ion coverage is annotated in blue, and the y-ion coverage in red.

**Figure 7.**
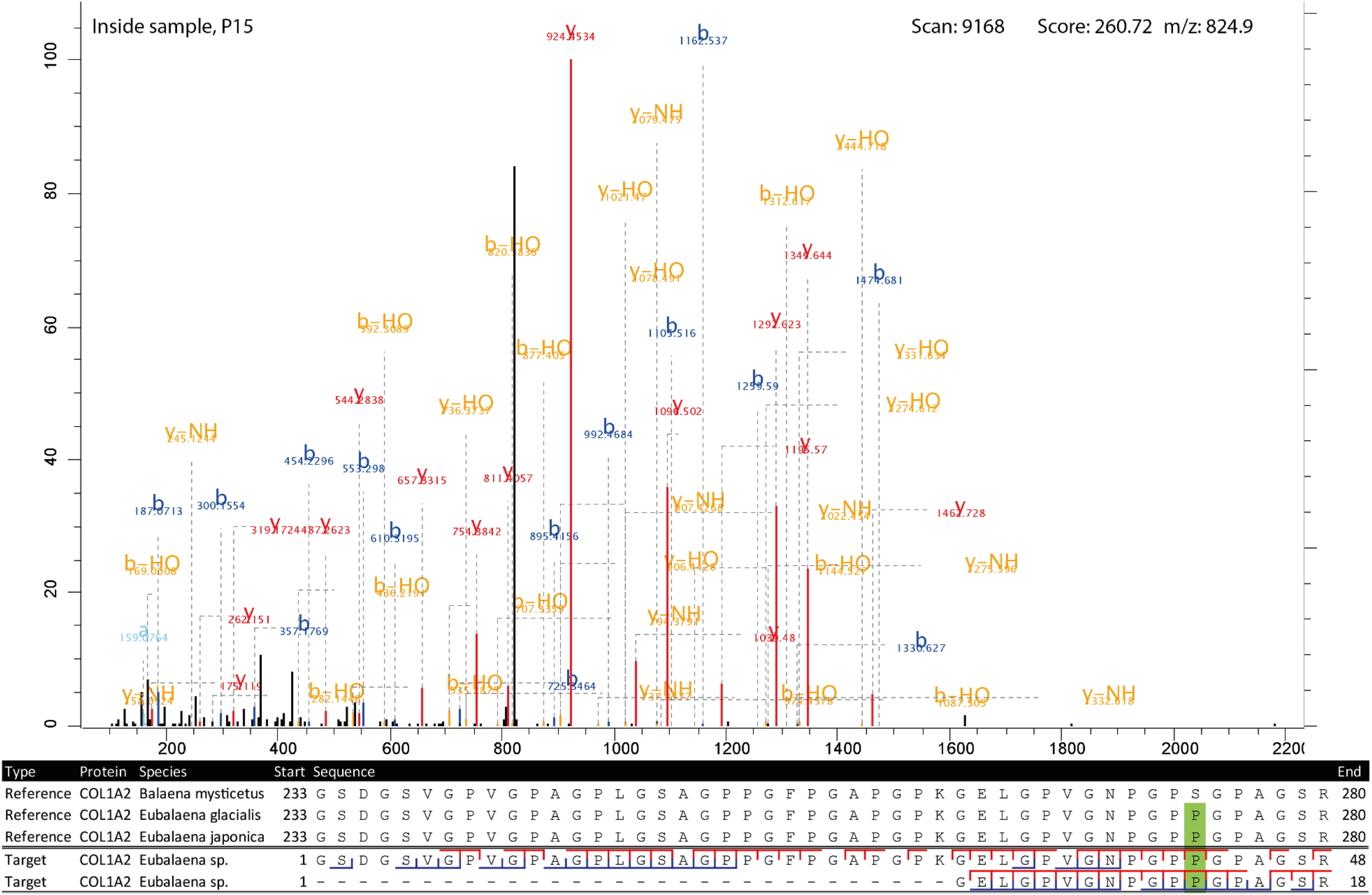
Highest scoring MS2 spectrum of the COL1A2 262-280 peptide covering the 276 SAP. The reference spectra matching the observed peptides are highlighted in green at the SAP. The b-ion coverage is annotated in blue, and the y-ion coverage in red.

## Discussion

### Contextualising molecular taxonomic identification

Three proteomic workflows have been applied in this study, each with different levels of specificity and requiring different approaches to critically interpret their results (Figure 8). A common feature is that in the initial phase of analysis no a priori assumptions on the presence or absence of species are made, but all species are considered as possibilities. The taxonomic identification should be made on a taxonomic level encompassing all species matching the proteomic signal, but excluding lower-level taxa with their own reference data that contradict the observed data. The baleen whale clade illustrates this principle well, as no reference data is available for several of its members, for both ZooMS and SPIN. Consequently, for example in the case of the ZooMS analysis of sample 2199, *Eubalaena australis* (southern right whale) could also be considered as a candidate, because no collagen biomarkers have been established for this species. It could be that the collagen sequence that sets *Eubalaena australis* apart from the *Eubalaena glacialis* and *Eubalaena japonica* sequences was ancestral to all members of Eubalaena. As a result, the other members of the Eubalaena genus for which we do not have reference biomarkers should be taken into consideration. Thus, to minimise the impact of limited reference data in proteomic identification of bone specimens, we constructed a database of collagen type I sequences capturing most of the existing sequence variation among extant baleen whales for the targeted database search. This database, which misses only *Balaenoptera edeni* and *Balaenoptera omurai*, could in turn be used to significantly expand the ZooMS peptide marker database. Additionally, the taxonomic resolution of the proteomic identifications can be greatly improved by filtering them based on the archaeological and geographical context of the sample.

**Figure 8.**
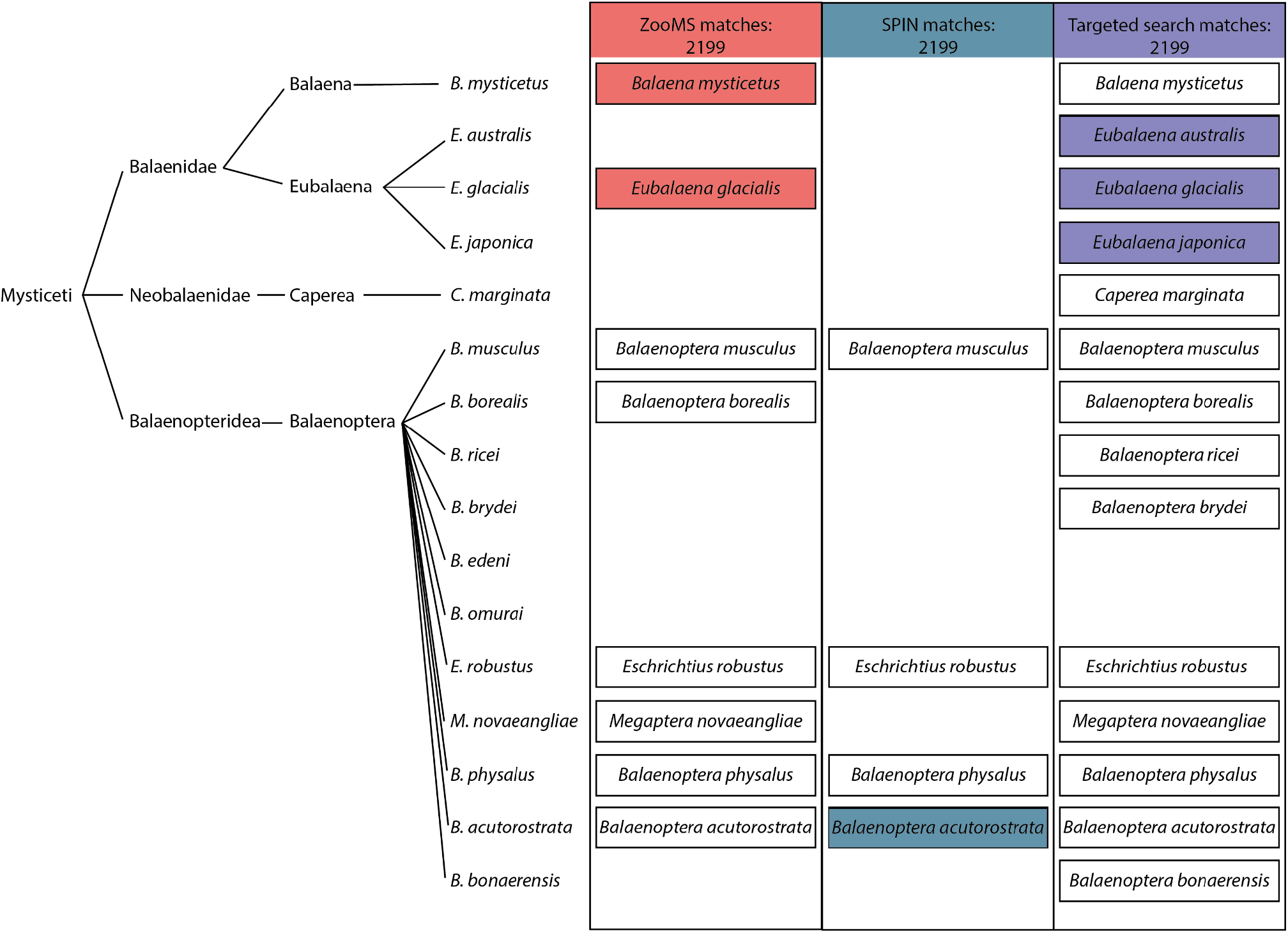
Baleen whale phylogeny and presence in relevant proteomic databases. Left: the phylogenetic tree of baleen whales based on recent genomic studies (Árnason et al. 2018; McGowen et al. 2020; Wolf et al. 2023). Right: tabular overview of the species that can be taxonomically identified using ZooMS and SPIN. Species for which reference biomarkers are available have a black border and those that match the observed biomarkers following their analytical workflow have a coloured background. Species absent for ZooMS, SPIN, and/or the targeted search are left out from each relevant column.

Here, we assess potential database biases and review the historical distributions of the possible species identified by the molecular analysis to exclude species that were either identified as false positives due to database limitations or not present in the North Sea during the Holocene (Figure 7). First of all, while the species is present in the North Sea, we argue that the *Balaenoptera acutorostrata* identification for the bone artefact specimen 2199 obtained by SPIN can be excluded from the possible identifications, because it is an artefact of the database selection. No Balaenidae were included in the SPIN database, whereas all except two baleen whale species were included in the database used in the targeted search. As both the ZooMS and targeted search analyses of specimen2199 used more comprehensive databases, including Balaenidae and Balaenopteridae references, the conflicting *Balaenoptera acutorostrata* identification can be safely regarded as a database artefact.

As for the remaining list of potential species candidates for specimen 2199, the archaeological context of Heiloo Zuiderloo allows the taxonomic identification to be narrowed down to a single species; *Eubalaena glacialis* (North Atlantic right whale). Ideally this identification would be based on securely identified remains from the same region and time period. However, due to the limited availability of species-level identified whales from Bronze Age Netherlands, we included archaeological findings from other periods, as well as the modern distribution of the candidate species. The presence of *Eubalaena glacialis* is attested in the Netherlands from the mid-Holocene (Foote et al 2013) and at least the first millennium AD onwards (van den Hurk et al. 2022) and it has been tentatively suggested that this species might have been one of the most frequently exploited cetaceans along the European Atlantic coast (van den Hurk et al. 2023). The two other candidate species, *Eubalaena japonica* (North Pacific right whale) and *Eubalaena australis* (southern right whale), are native to the North Pacific Ocean (Kenney 2009) and the southern hemisphere, respectively (Richards 2009) and can therefore be safely excluded as potential sources of sample 2199. Instead, it seems most likely that the adze was produced from a bone of *Eubalaena glacialis*. The proteomic taxonomic identification of the two objects deviates substantially from the original morphological estimation. However, it seems that for fragmented remains it can be particularly difficult to distinguish Elephantidae and Mysticeti. In fact, several cases of biomolecular techniques unmasking morphologically identified Elephantidae remains are known (van den Hurk et al. 2020), highlighting that this may be an area where accessible biomolecular taxonomic identifications, such as ZooMS, are particularly useful.

### Bronze Age whale exploitation: hunting or scavenging?

Prehistoric baleen whale remains were until recently considered rare in the North Sea region, but are increasingly reported. Aaris-Sørensen et al. (2010) identify at least 100 baleen whale Pleistocene and Holocene remains in Denmark with several dating to the Bronze Age, and Foote et al. (2013) documented the presence of North Atlantic right whales and bowhead whales in the Netherlands back to the early Holocene and Pleistocene, respectively. These finds make it clear that the North Sea region has a long history of baleen whale exploitation, but whether this consisted of the opportunistic use of beached whales or active hunting remains a question for a large part of this history. It is impossible to determine if a single bone is derived from a beached or hunted animal, but several characteristics have been proposed that would suggest it is more likely that a whale bone assemblage was obtained by active whaling rather than opportunistic scavenging. Active whaling assemblages are thought to be relatively limited in taxonomic diversity (van den Hurk et al. 2023; Wellman et al. 2017), whale remains must be abundant and there must be some form of industry associated with the whale remains, such as bone artefact production or blubber processing (Hennius et al. 2018, 2023; Wellman et al. 2017). A potential side-effect of this requirement is that active whaling purely for consumption becomes more difficult to detect archaeologically than whaling to acquire resources for product manufacture. Lastly, written sources are frequently used as evidence for active whaling in historical periods (Hennius et al. 2018; van den Hurk et al. 2023), but these are not available for prehistoric periods. In general, most studies on active whaling agree that it needs to be accompanied by a certain scale of whale resource exploitation.

Early suggested examples of active whaling are the Dutch Neolithic Vlaardingen Culture (van den Hurk et al. 2023) and 6th century Scandinavia (Hennius et al. 2018). Then from the Middle Ages onwards active whaling seems to become more prevalent along the Atlantic coast, most notable are the Basques and northern Spaniards starting from at least the 11th century (Rey-Iglesia et al. 2018), but also in the Netherlands (van den Hurk, Spindler, and McGrath 2022). The evidence for active whaling in the Vlaardingen Culture consists of 13 and 28 baleen whale bones at two sites respectively, 10 and 7 of which were identified as *Eschrichtius robustus* (the grey whale). The dominance of *Eschrichthius robustus* suggests a taxonomic focus characteristic of active whaling.

Additionally, the sites indicate an overall marine focus of the culture, displayed by their diet and material culture (Brinkkemper, Drenth, and Zeiler 2011; van den Hurk et al. 2023). The case for active whaling in 6th century Scandinavia is more robust. The large increase in the production of whale bone gaming pieces across Scandinavia, as well as an increase in blubber processing pits indicates a significant increase in the exploitation of whale resources (Hennius et al. 2018). Additionally, most of the whale bone species were assigned to *Balaenidae* sp., again suggesting a hunting preference for a particular taxon (Hennius et al. 2023). For the mediaeval active whaling cultures of the Basques and Northern Spaniards and the Dutch there are also historical sources attesting to active whaling next to the archaeological bone assemblage. Both these cultures also seem to have favoured *Eubalaena glacialis* as their target for whaling (Rey-Iglesia et al. 2018; van den Hurk et al. 2022).

The aforementioned examples of active whaling can be used to evaluate the baleen whale presence in the archaeological record of the Dutch Bronze Age against. Apart from the *Eubalaena glacialis* artefact described in this study, eight other cetacean remains have been found at the same site of Heiloo Zuiderloo (Heiden 2018; de Koning and Tuinman 2019; Moesker et al. 2021), one of which was also identified as Balaenidae and two others as grey whales (van den Hurk et al. 2023). Additionally, several Bronze Age whale remains have been found relatively close to Heiloo. Two whale ribs dated to 1550-1350 BC have been found near Bergen (de Ridder 1993), another whale rib was found in Boekelermeer, dated to around 1500 BC, and two whale bones were found in Alkmaar, one being dated to the Middle Bronze Age (Kleijne 2015). The ribs from Bergen and Boekelermeer have recently been identified using ZooMS as sperm whale (*Physeter macrocephalus*) and grey whale (*Eschrichtius robustus*) respectively (van den Hurk et al. 2023). The remains from Alkmaar could not be identified beyond ‘whale’, which likely is supposed to refer to any large-bodied cetacean.

Regarding whether these cetacean remains are the result of active whaling or scavenging, in the case of the Bergen ribs they were said to have been found in a marine deposit. The authors hypothesise that these two ribs are the remains of a stranded whale butchered on the beach where it landed (de Ridder 1993). Considering the remaining finds, the evidence for active whaling does not appear strong. Although there is a cluster of cetaceans from a relatively narrow time period in a small area, they feature a high taxonomic diversity more indicative of opportunistic exploitation of beached whales. We cannot exclude active whaling, but neither does there appear to be strong evidence for the presence of active whaling in the Dutch Bronze Age.

### Contextualising the tool in the Bronze Age

The current study has mainly been devoted to studying the first step in a bone tool biography: the choice of the raw material for the tool. However, the find location of the tool raises questions regarding the last chapter in its biography: its deposition, particularly the intentionality behind the deposition. The object was found near a local depression in the dune landscape. Extensive studies have shown that in Bronze Age Europe, these natural wet locations were desirable places for intentional deposition and ’destruction’ of what we consider valuable objects (Fontijn 2002, 2020). In the vicinity of the 2020 excavation location, objects associated with such a phenomenon have been discovered in the past. These include a group of four Late Bronze Age flint sickles and a bronze sickle together inserted upside down in the peat (Brunsting 1962) and an unused Middle Bronze Age stop-ridge axe that was probably made in Normandy (Moesker et al. 2021). Although the Heiloo specimen was made of an ‘exotic’ material, use-wear traces show that it has been extensively worked and used. Efforts have been made to shape and finish the object and long-term use has left traces on both the bone part and the wooden handle of the object. Although the artefact is not broken and does not appear to have been irreparably damaged, it ultimately ended up in an active watering hole that was emptied at a later time. The evidence suggests that it is not a deliberate deposition, but rather a lost object, although the first cannot be ruled out.

The adze-like tool may be lost in more ways than one. No parallel of the object is currently known. Tools made of bone or antler with a distinct working edge are known from many locations in the Netherlands during the Bronze Age (Zeiler et al., forthcoming). Although no clear parallel for this object can be found, it is notable that in recent years, several other utensils associated with the processing of organic materials such as flax and fibres have been found in Heiloo (Tuinman et al. 2022; Koning and Tuinman 2021; Lange 2021; Edmonds 2018; Verbaas and Edmonds 2018). These involve tools made of organic materials, including a bone comb and two oak wood pestles. It demonstrates the significant importance of the prehistoric sites in Zuiderloo and the unique preservation conditions in which such organic tools are preserved, but at the moment it does not provide much clarity regarding the intended purpose of the tool nor its place within the Bronze Age toolkit.

## Conclusion

Although in recent years advances in biomolecular applications to archaeology have facilitated the taxonomic identification of osseous tools, there remain some taxonomic and methodological challenges. This small scale study on an exceptional bone artefact from Bronze Age Heiloo exemplifies well how biomolecular methods can improve our understanding of what species were exploited, as well as the different challenges and pitfalls that it entails. After performing ZooMS, SPIN, and shotgun-proteomic data analysis and while considering their individual methodological strengths and limitations, we have shown that the extraordinary adze-like bone tool was most likely manufactured from the bone of *Eubalaena glacialis* (North Atlantic right whale). Use-wear analysis furthermore demonstrates that it was used for the processing of plant fibres. We cannot be certain whether the bone to produce the artefact was obtained from a beached whale or by active whaling, but our current understanding of cetacean remains in the Dutch Bronze Age does not point towards the existence of an active whaling practice. Additionally, we contribute previously unavailable COL1A1 and COL1A2 reference sequences for six baleen whale species, in order to facilitate the future proteomic identification of these species within the archaeological and palaeontological record.

## Acknowledgements

We would like to thank André Ramcharan and Dr. Laura Llorente Rodriguez for their initial morphological identification and Prof. Marie Soressi for her advice on setting up this project. Special thanks go to Femke Reidsma, Erica van Hees, Dr. Rachel Schats and Dr. Jason Laffoon for their aid in sampling the adze. We thank the IZI Fraunhofer, S. Kalkhof and J. Schmidt for providing access to the MALDI–TOF MS instrument in Leipzig, Germany and the York Centre of Excellence in Mass Spectrometry for the use of the Ultraflex III MALDI-ToF/ToF instrument. Lastly, we would also like to express our gratitude to Meaghan Mackie for her work in acquiring the LC-MS/MS data.

## Funding

J.D. is funded by a Marie Skłodowska-Curie grant (grant agreement No. 956351) from the EU Horizon 2020 programme.

This research has been made possible through funding from the European Research Council (ERC) under the European Union’s Horizon 2020 research and innovation programme, grant agreement no. 948365 (PROSPER) awarded to F.W.

D.M. received support from the European Union’s Horizon 2020 research and innovation programme under the Marie Skłodowska-Curie grant agreement no. 861389 (PUSHH).

M.L.M. is funded by a Carlsberg Foundation Semper Ardens Accelerate Grant CF21-0425, awarded to M.T.O.

V.S.-M. is supported by a Fyssen Foundation postdoctoral fellowship (2023-2025). ZooMS analyses performed in Leipzig were financially supported by the Max Planck Society.

The excavation at Heiloo-Zuiderloo and subsequent analysis by J.B. were performed as part of the commercial activities of Archol BV commissioned by the municipality of Heiloo. The use-wear analysis by A.V. in turn was commissioned by Archol BV.

## Conflict of interest disclosure

The authors declare that they comply with the PCI rule of having no financial conflicts of interest in relation to the content of the article.

Data, scripts, code, and supplementary information availability (if applicable) The MALDI-MS spectra used for ZooMS can be found on Zenodo with the following identifier: https://zenodo.org/doi/10.5281/zenodo.10970628.

The LC-MS/MS data was deposited on the ProteomeXchange Consortium via the PRIDE partner repository under the dataset identifier PXD051408.

